# The Role of Feedforward and Feedback Inhibition in Modulating Theta-Gamma Cross-Frequency Interactions

**DOI:** 10.1101/2025.01.24.634658

**Authors:** Dimitrios Chalkiadakis, Jaime Sánchez-Claros, Víctor J López-Madrona, Santiago Canals, Claudio R. Mirasso

**Author notes:** Correspondence: Claudio R. Mirasso, Santiago Canals.

## Abstract

Interactions among brain rhythms play a crucial role in organizing neuronal firing sequences during specific cognitive functions. In memory formation, the coupling between the phase of the theta rhythm and the amplitude of gamma oscillations has been extensively studied in the hippocampus. Prevailing perspectives suggest that the phase of the slower oscillation modulates the fast activity. However, recent metrics, such as Cross-Frequency Directionality (CFD), indicate that these electrophysiological interactions can be bidirectional. Using a computational model, we demonstrate that feedforward inhibition modeled by a theta-modulated ING (Interneuron Network Gamma) mechanism induces fast-to-slow interactions, while feedback inhibition through a PING (Pyramidal Interneuron Network Gamma) model drives slow-to-fast interactions. Importantly, in circuits combining both feedforward and feedback motifs, as commonly found experimentally, directionality is flexibly modulated by synaptic strength within biologically realistic ranges. A signature of this interaction is that fast-to-slow dominance in feedforward motifs is associated with gamma oscillations of higher frequency, and *vice versa*. Using previously acquired electrophysiological data from the hippocampus of rats freely navigating in a familiar environment or in a novel one, we show that CFD is dynamically regulated and linked to the frequency of the gamma band, as predicted by the model. Finally, the model attributes each theta-gamma interaction scheme, determined by the balance between feedforward and feedback inhibition, to distinct modes of information transmission and integration, adding computational flexibility. Our results offer a plausible neurobiological interpretation for cross-frequency directionality measurements associated with the activation of different underlying motifs that serve distinct computational needs.

**Author Summary:** This study investigates the interaction between various types of brain oscillations and their potential relationship with the connectivity of underlying neural networks. Brain activity encompasses slow oscillations, such as theta, alpha, and delta, as well as faster oscillations, including gamma. These oscillations interact through Cross-Frequency Coupling (CFC), a mechanism essential for cognitive processes like memory, learning, and attention. Given the higher spectral power and broader spatial propagation of slow oscillations, it has been proposed that CFC arises when slow oscillations modulate faster activity. However, recent evidence suggests that gamma oscillations can also predict the phase of slower oscillations, indicating a bidirectional and more intricate relationship. To explore this complexity, we developed a computational model that reproduces both forms of interaction observed experimentally. Our results demonstrate that while slow oscillations originating from distant regions can induce gamma activity, local connectivity and specific cell-type dynamics allow gamma oscillations to anticipate slow oscillations in certain conditions. The balance of inhibitory circuits modulates fast-slow oscillation interactions, creating distinct dynamical modes with varying computational properties and enhancing system flexibility. This work integrates competing hypotheses on oscillation interactions and offers a conceptual framework for linking these dynamics to the structural organization of neural circuits.

## Introduction

Mammalian brains exhibit oscillatory activity over a broad frequency range, from 0.5 to 500 Hz, spatially organized across different regions [1]. These oscillations reflect distinct synchronization patterns of the underlying neural circuits and are associated with different behavioral states. For instance, theta band oscillations, around (4-8) Hz, in the prefrontal cortex and hippocampus and alpha band oscillations, around (8-12) Hz, in the visual cortex, have been linked with locomotion, learning, and attention [2–6].

Faster rhythms, such as gamma oscillations within the 30-150 Hz frequency band, are also ubiquitous in brain networks [7, 8]. Notably, gamma oscillations frequently interact with slower rhythms in a phenomenon known as cross-frequency coupling (CFC) [9]. The hippocampus in particular, has been extensively studied for its theta-gamma CFC, which shows increased gamma amplitude locking to the phase of the theta cycle during decision-making and learning [10–12]. Additionally, CFC has been observed in the cortex, where gamma activity couples with theta, alpha, and beta oscillations [13]. Theories of neural computation propose that CFC plays a critical role in inter-regional communication essential for attention [14], and in organizing neuronal firing into cell assembly sequences underlying episodic memory formation [15].

The generation of CFC, though not fully understood, has been explained based on local and global network properties. *In vitro* and computational studies identified intrinsic neuronal properties, such as I_h_ currents, and interactions between fast and slow interneurons, as key local mechanisms [16–20]. Simultaneous electrophysiological recordings in the hippocampus and entorhinal cortex showed, however, that CFC is also influenced by rhythmic inputs from upstream layers [21–23]. The finding of high coherence at low frequencies across distant recording sites, but not at high frequencies, led to the concept that slow oscillations are driven by upstream areas, which then locally organize faster network dynamics [24–26]. This aligns with the oscillatory hierarchy hypothesis, which posits that slower oscillations modulate population excitability, thereby coordinating higher-frequency activity [27].

However, the application of techniques for separating field potentials into pathway- or layer-specific activity patterns- [28], combined with new metrics that assess directional interactions across frequency bands, such as cross-frequency-directionality (CFD) [29], has revealed bidirectional interactions between fast and slow oscillations. In the hippocampus, recordings from rats engaged in navigation and memory tasks demonstrated that bouts of gamma activity systematically preceded the phase of theta oscillations, suggesting gamma-to-theta interaction [12] (see Fig. 1a). Similarly, human electrocorticography studies have reported gamma-to-alpha interactions in the visual cortex [29, 30]. Furthermore, using two independent methods to assess directionality, Dupré la Tour and colleagues [31] observed both interactions: theta-to-gamma in the hippocampus during REM sleep in rats, and gamma-to-theta in the human auditory cortex. Overall, CFC is not exclusively due to slow-to-fast interactions challenging the conventional oscillatory hierarchy hypothesis.

**Fig. 1.**
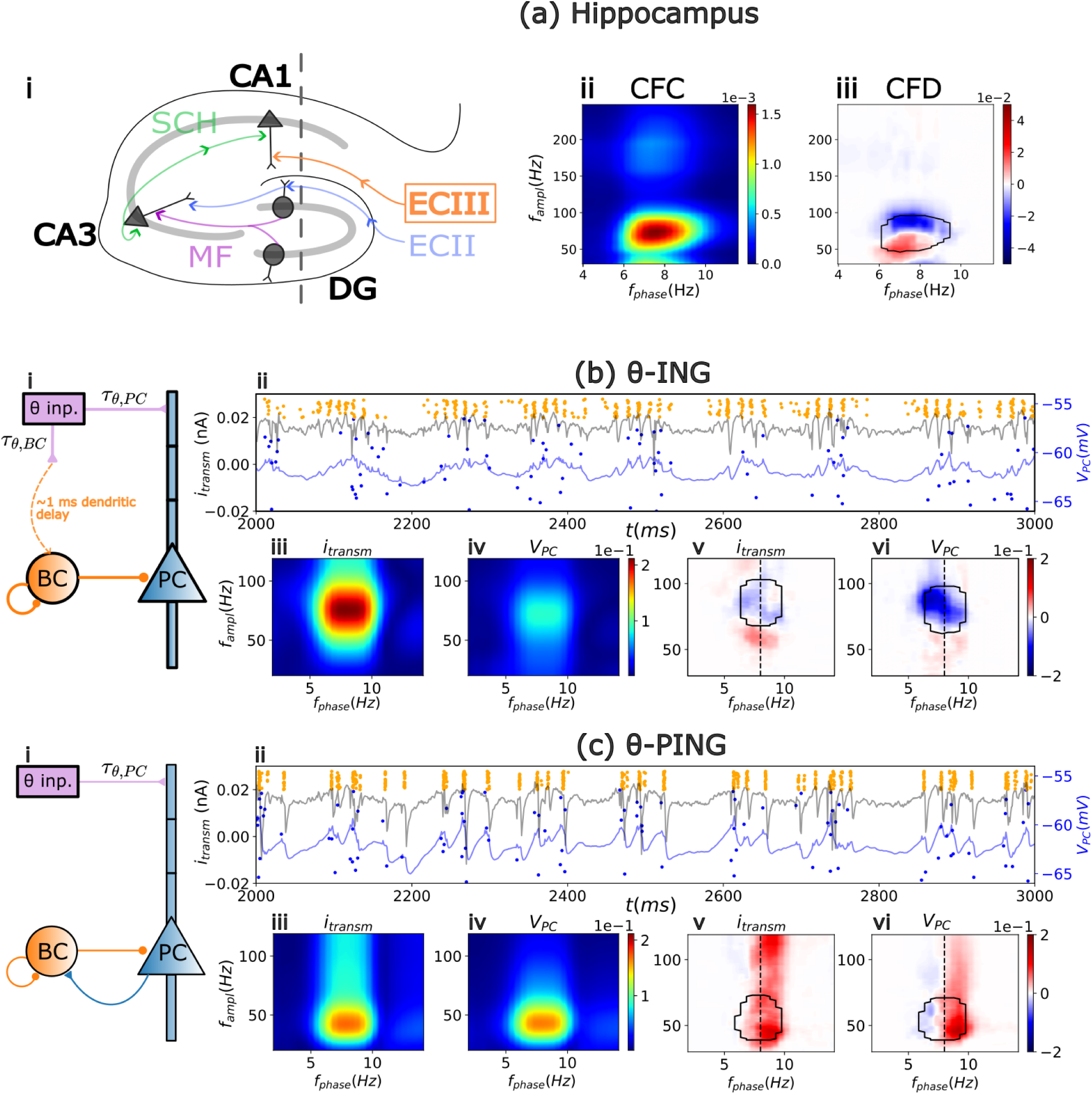
Experimental and computational cross frequency coupling and directionality. (a) Cross-Frequency Coupling (CFC) and Cross-Frequency Directionality (CFD) obtained from electrophysiological recordings from the rat hippocampus (data from [12]). (i) Schematic representation of the hippocampus with the pathway-specific layers illustrated. Sch: Schaffer Collateral, MF: Mossy Fibers, ECII and ECIII: Entorhinal Cortex layers II and III, DG: Dentate Gyrus, CA1 and CA3: Cornu Ammonis. (a-ii) Theta-gamma CFC in the field potential associated with the ECIII and (a-iii) its corresponding CFD. Note that averaging the CFD over areas of high CFC (black contour in CFD panels) shows a predominantly negative directionality CFD_avg_ = -0.014. (b) θ-ING model. (i) Circuit motif. (ii) Timeseries of the average PC somatic transmembrane current (i_transm_), average somatic PC membrane voltage (V_PC_) and raster plots of the BCs (orange dots) and PCs (blue dots) firing. The corresponding CFC and CFD for the i_transm_ and V_PC_ are depicted in panels (iii and v) and (iv and vi), respectively. The black contour in CFD plots shows the 90 percentile of the respective CFC. (c) Same as (b) but for the θ-PING motif.

In this study, we aim to (1) address and reconcile this discrepancy by employing a computational modeling approach and (2) offer a plausible neurobiological interpretation of CFD. Specifically, we adapt a model from the work of [32] to investigate the underlying mechanisms and provide new insights. In our model, pyramidal neurons receive inputs from an external population generating a slow (theta) rhythm, while the fast (gamma) rhythm emerges locally through the activity of fast-spiking interneurons. We found that the local cross-frequency directionality is determined by the dominance of specific connectivity motifs within the underlying circuit. In a theta-modulated Interneuron Network Gamma (θ-ING) motif (Fig. 1b), feedforward recruitment of fast-responding interneurons primarily drives gamma-to-theta directionality. In contrast, in theta-modulated Pyramidal-Interneuron Network Gamma (θ-PING) motif (Fig. 1c), feedback inhibition supports theta-to-gamma directionality. In combined motifs that reflect the anatomy of neuronal circuits commonly found in the brain, we analyzed transitions between these modes, demonstrating smooth bidirectional interactions controlled by synaptic strength within biologically plausible ranges. We validated several model predictions using previously acquired experimental data [12]. Finally, by evaluating the capacity of each computational motif to integrate distinct inputs, we uncovered a mechanism that prioritizes transmission across different parallel information channels arriving at the dendrites of pyramidal cells.

## Methods

### Theta-gamma generation in the θ-ING and θ-PING motifs

The two motifs analyzed in this study are variations of the Interneurons Network Gamma (ING) and Pyramidal-Interneuron Network Gamma (PING) models, both of which generate gamma activity through interactions between pyramidal cells (PCs) and fast-spiking inhibitory interneurons, primarily parvalbumin-immunoreactive basket cells (BCs) [33–37]. In both models, a gamma cycle begins with BCs firing, which suppresses the activity of their target neurons (both PCs and other BCs). Once the inhibition decays, BCs fire again, initiating a new cycle. The key difference between the two models lies in the source of excitatory drive to BCs: in the ING model, excitation originates from outside the network, whereas in the PING model, it results from bidirectional interactions between BCs and PCs.

We modified the traditional ING and PING models by incorporating an external theta drive that regulated the activation and suppression of gamma dynamics (Fig. 1b and Fig. 1c), thereby inducing theta-gamma cross-frequency coupling (CFC). The θ input was modeled such that the interval between consecutive peak activations, T_cycle_, was drawn from a Gaussian distribution with a mean μ_T_ = 125ms and a standard deviation σ_T_ = 16ms. Each cycle contained 10,000 spikes, distributed according to a second Gaussian with a standard deviation *w* = 25ms. The variability in both inter- and intra-cycle timing, controlled by σ_T_ and *w*, respectively, was essential to detect directional interaction between theta and gamma rhythms.

In order to make meaningful comparisons between the two motifs we used the same synaptic weights for the connections θ→PCs and BCs→PCs while the PCs→BCs synaptic strength of the θ-PING was chosen so that the mean firing rate of the PCs was almost identical in both motifs (see Table S2 for more details). For instance, in Fig. 1 the firing rates are f_r,θ-ING_ = 0.48(0.01) Hz and f_r,θ-PING_ = 0.49(0.01) Hz, for the mean value and the standard deviation (in parenthessis) across 20 simulations. Nonetheless, the gamma oscillations of the whole population differed in the θ-ING and θ-PING motifs, with the θ-ING one exhibiting faster dynamics (Fig. 1b-ii,iv vs 1c-ii,iv). This occured because the θ input’s strength was set to a relatively low value, causing sparse firing of the individual neurons at a rate lower than the population frequency, similar to what is observed in vivo. This discrepancy was even more pronounced in PCs, to the extent that computational models replicating this behavior are classified within the weak PING/ING subcategory [38]. Thus, due to their lower input and slower dynamics, PCs required more time to activate BCs in the θ-PING motif compared to BCs in the θ-ING model, which were directly driven by the θ input.

### Cells

Although the motifs we analyzed are ubiquitous in multiple brain areas, our model was adapted from previous works modeling the hippocampus CA3 area in [32]. Pyramidal cells were represented as multi-compartmental neurons with active dendrites while basket cells were modeled as single-compartment point neurons. Both neuron types included leak, transient sodium, and delayed rectifier potassium currents, with PCs also incorporating hyperpolarization-activated currents. For a comprehensive description of the model see also chapter 4 section 4.2 of [39] and ref. [32]; the simulations code is available in GitHub (https://github.com/gerompampastrumf/thetaING_PING).

Modeling multicompartmental pyramidal cells with dendrites allows for the simulation of realistic transmembrane currents and Local Field Potential (LFP) recordings [40]. To achieve this, our PCs were designed with five compartments, each subdivided into three segments to enhance spatial integration accuracy. Panels (b-i) and (c-i) of Fig. 1 depict a scheme of the model including the morphology of the PC. 3 compartments emulated the apical dendrites, one the soma and another one the basal dendrites. All compartments were modeled as cylinders.

For the BCs we chose not to model the dendritic tree explicitly but considered them as point neurons, following ref. [32]. However, since dendritic transmission time in BCs is small but not negligible, we incorporated an additional fixed delay of 1 ms from the external driver to the basket cells. This dendritic delay was selected to exceed the onset time between evoked dendritic postsynaptic currents and the corresponding increase in somatic membrane potential observed in patch-clamp experiments (see Fig. 3d of [41]).

### Synapses

Our model included three types of synaptic connections: inhibitory GABA_A_, excitatory AMPA and NMDA, all of which were modeled using the standard double exponential function [32]. Synaptic weights for NMDA connections were fixed at a value ten times lower than those for AMPA. Neuronal connections were chosen randomly. Axonal propagation and synaptic transmission delays between populations *x* and *y* were modelled using a transmission delay τ_x,y_, drawn from a Gaussian distribution with a standard deviation of 0.2 ms. The mean of the distributions were set as follows: τ_θ,BC_ = 20 ms, τ_θ,PC_ = 20 ms, τ_BC,PC_ = 1.5 ms, τ_PC,BC_ = 1.5 ms, and τ_BC,BC_ = 1.5 ms. The relatively large values for τ_θ,BC_ and τ_θ,PC_ were chosen to ensure that, in Fig. 2, the relative difference between the two could be manipulated without resulting in negative delays. To account for background activity, poissonian noise sources with a rate of 1 ms were added to the pyramidal soma, proximal dendrites, and basket cells. In the simulations shown in Fig. 5, we examined the competition between a parallel pathway, incorporated in the proximal dendrite, and the theta drive. The parallel pathway was modeled as a Poissonian source with its synaptic strength increased tenfold with respect to the background activity and its rate adjusted from 1 ms to 5 ms. The values of the synaptic parameters are detailed in Tables S1 and S2 in the supplementary material section.

**Fig. 2.**
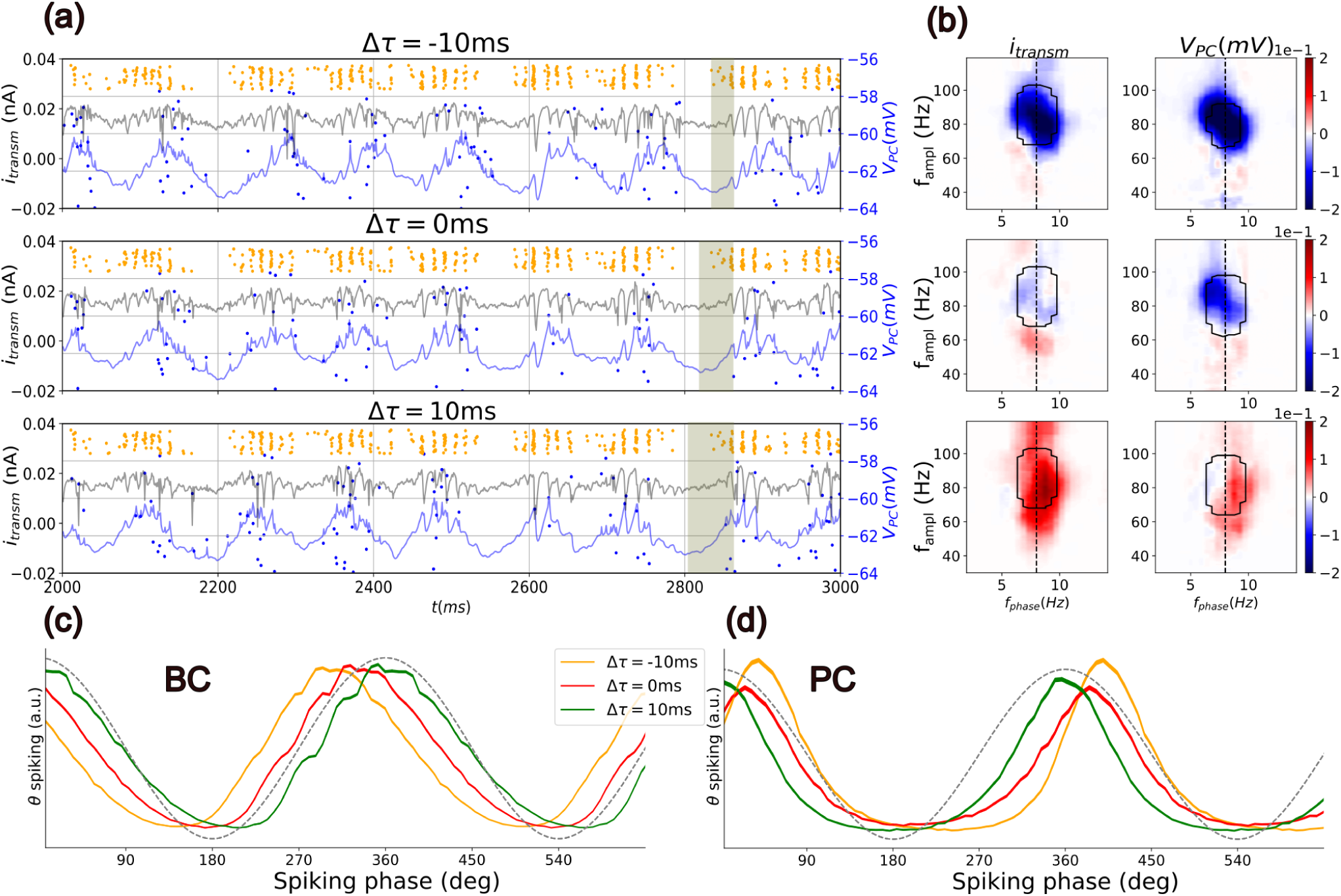
Directionality in the θ-ING model reverses depending on the relative transmission delay. (a) Timeseries of i_transm_ (black line) and V_PC_ (blue line) and raster plots of the BCs (orange dots) and PCs (blue dots) spikes, for different relative transmission delay Δτ. The shaded area highlights the relative displacement of the theta trough (minima of i_transm_) with respect to the initiation of the gamma oscillation. (b) CFDs for i_transm_ and V_PC_. Contour lines indicate the range of higher CFC (more than the 90th percentile). Relative transmission delay Δτ = -10/0/10 ms increases from top to bottom in (a) and (b). In panels (c-d) we plot the theta-phase spiking distributions for BCs and PCs, respectively. For this purpose, the synaptic current generated by the θ input (indicated by the dashed black line) was used as a reference signal to extract a theta phase. The spiking profiles of the BC and PC was then calculated by assigning the corresponding phase to each spike.

**Fig. 3.**
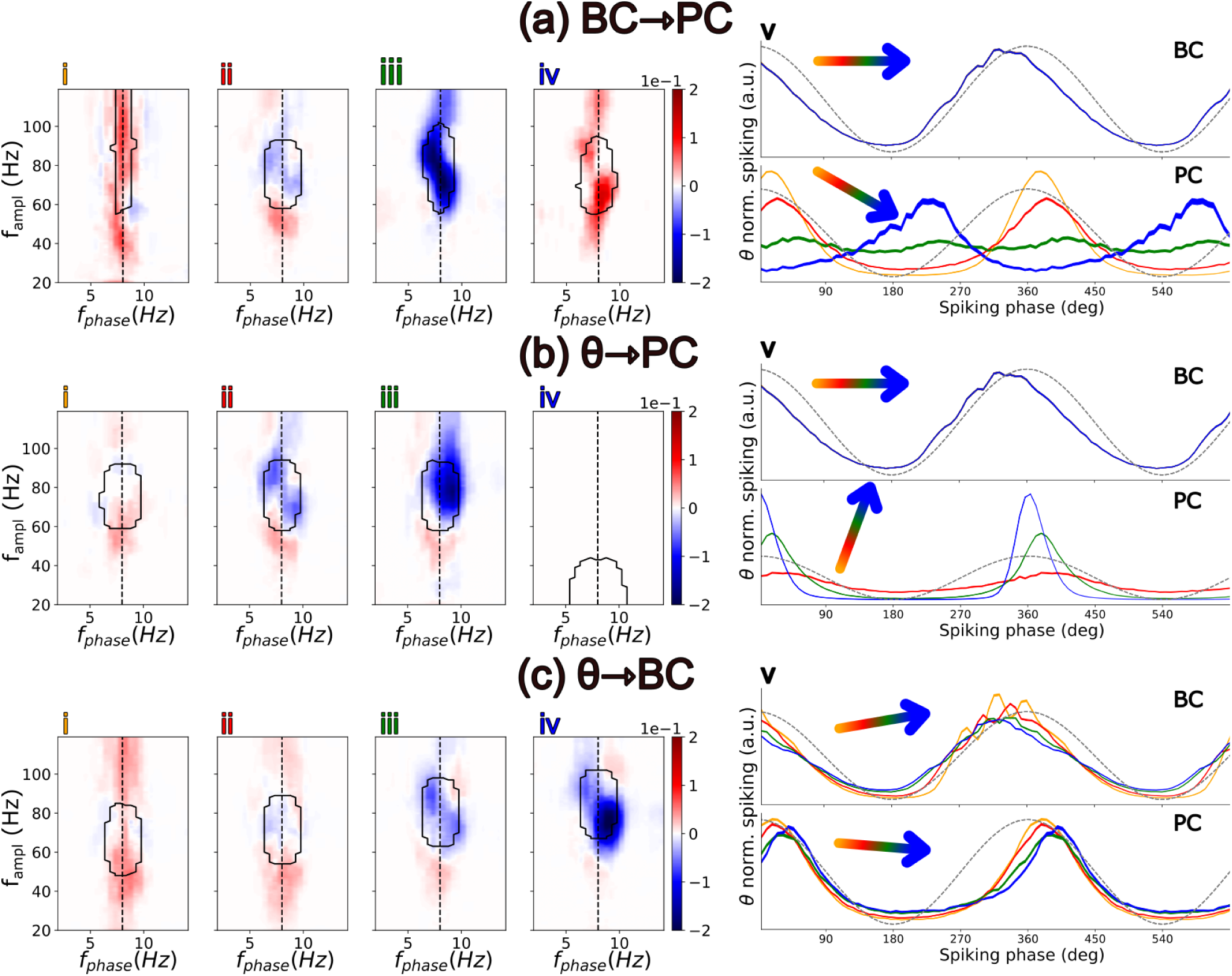
Role of different synaptic coupling strengths on directionality, firing rates, and spiking phase. The synaptic coupling takes four increasing values: w_i_<w_ii_<w_iii_<w_iv_ (see Table S3 for exact values) for the following connections: BC→PC (a), θ→PC (b), and θ→BC (c). (i-iv) CFD with the contour encircling areas of high CFC values. (v) Spiking distribution as a function of the theta phase for BCs (Top) and PCs (Bottom). Spikes are binned according to the phase of the synaptic current of the θ input in the PCs dendrites (same as in Fig. 2). For visualisation purposes, each spiking distribution was normalized so that the integral over a theta cycle is 1. The orientation angle of the coloured arrow indicates whether the firing rate increases, decreases, or remains the same as the corresponding synaptic weight increases (see Table S4 for exact values).

### Simulations

Simulations were conducted using the NEURON simulator library [42]. Each simulation modeled 60 seconds of activity across 20 realizations, with different realizations of the Poissonian and θ-inputs. The only exception was the simulations used to generate Fig. 5, where, depending on the protocol, a selected input was kept fixed across simulations, while different realizations were used for the remaining inputs. Each simulation included 200 PCs and 40 BCs.

### Dataset

All experimental data used in this study were previously acquired and described in [12]. Briefly, the dataset (available at https://doi.org/10.20350/digitalCSIC/12537) comprises electrophysiological recordings (32-channel silicon probes) from five rats along the dorso-ventral axis of the dorsal hippocampus, covering the CA1 region and the dentate gyrus. During the recordings, the animals freely explored an open field to which they had been previously habituated over the course of one week through daily sessions (familiar environment condition). Subsequently, a modification was introduced to the open field—a change in the surface texture of the floor—and the animals were recorded again (mismatched novelty condition). The results of theta-gamma CFC and CFD under these conditions can be found in [12]. The specific dataset used in the present study consists of pathway-specific field potentials from the lacunosum-moleculare layer of CA1, which exhibits the highest signal-to-noise ratio for theta and gamma activity. These field potentials were obtained by applying Independent Component Analysis (ICA) to the 32 time series of local field potentials (LFP) recorded using the multichannel electrodes [28, 12].

### Analysis

#### Cross Frequency Coupling

This metric aims to quantify the degree to which the amplitude of a signal at a given high frequency f_ampl_ co-modulates with the phase at a different low frequency f_phase_. Various metrics have been developed for this purpose, but we will use the one proposed by Canolty et al. in [43], also known as the mean vector length. This method involves first filtering the signal at both f_ampl_ and f_phase_, followed by the calculation of the analytic signal’s amplitude at high frequencies Α_ampl_(t) and the analytic signal’s phase at low frequencies φ_phase_(t). Next, we construct the composite signal z(t) = Α_ampl_(t)exp(iφ_phase_(t)), which resides in the complex plane. If there is no coupling between the selected frequencies, the trajectory of z(t) will be radially symmetric. Consequently, the absolute average of the composite signal, z_avg_, is zero when there is no CFC and positive otherwise. In this study, we used the Comodulogram class from the pactools Python library created by Dupré la Tour and colleagues [31] to extract the absolute value of z_avg_.

To statistically assess the significance of the results, we employ a surrogate analysis followed by clustering correction for multiple comparisons [29]. We generate 1,000 surrogate time series by randomly splitting the phase signal into two segments and swapping their order. This procedure disrupts the temporal relationship between φ_low_(t) and Α_high_(t) while preserving key characteristics such as their spectra. We then calculate the 99th percentile value, (k_th_), which represents the CFC value greater than 99% of the samples across all surrogate data. All CFC values below k_th_ are set to zero. Subsequently, clusters of adjacent non-zero CFC values are identified within each dataset, both surrogate and non-surrogate. Each cluster is assigned a cluster score calculated as the sum of the values within the cluster. The CFC of the original data is considered significant at p<0.01 if its cluster score exceeds the 99th percentile of all surrogate cluster scores.

#### Cross Frequency Directionality

While the mean vector length metric effectively detects co-modulation of slow and fast components within a signal, it does not provide insights into potential temporal relationships between them. To address this limitation, we use the cross-frequency directionality (CFD) metric developed in [29]. CFD utilizes the phase slope index (PSI), which quantifies the directionality between two broadband signals, *x* and *y*, by analyzing the relationship of their phase difference, Δφ = φ_x_-φ_y_ as a function of frequency. When the phase differences are linearly dependent on frequency, it indicates a fixed time lag between the two signals. Specifically, if Δφ increases with frequency, *x* leads *y*; conversely, if Δφ decreases, *y* leads *x*.

In more detail, the PSI is calculated as follows. First, we compute the Fourier Transform of the signals *x* and *y* denoted *X* and *Y*, respectively. These series are then split into N smaller segments and the complex coherence is computed:

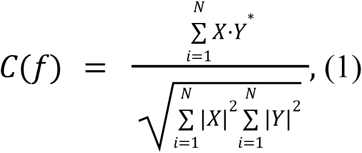

where ‘*” denotes the complex conjugate. Then, the phase slope index is calculated as:

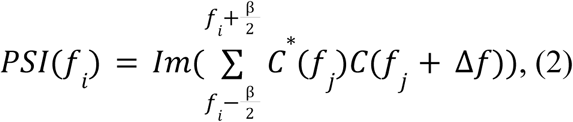

where “Im” refers to imaginary part. In cross-frequency analysis, the PSI is adapted when *y* is not a separate signal from *x*, but rather its amplitude filtered around a high frequency, f_high_. Finally, to emphasize the directionality of regions exhibiting strong cross-frequency coupling, we apply the masking technique described in [12]. Specifically, we devided the CFC by its maximum value so that it is bounded between 0 and 1, thus creating a mask. The final CFD is then calculated as the phase slope index multiplied by this mask.

To statistically assess the significance of the results, we employ the same procedure as for CFC for the highest 99th quantile. We also repeat the process for values lower than the 1st quantile in order to detect regions of significant negative CFD [29].

In certain cases, such as in Fig. 1a and Fig. 5, the predominant CFD_avg_ is extracted from the two-dimensional comodulogram. To achieve this, we generate an alternative binary mask over the CFC, retaining only frequencies where CFC exceeds the 0.95 quantile. We then compute CFD_avg_ by averaging all CFD values at these selected frequencies. Furthermore, we confirm that the results remain consistent when using alternative quantiles, such as 0.9 and 0.85.

#### From spikes to continuous timeseries

As part of the analysis, we examined the instantaneous output of a neuronal population. To achieve this, discrete spike events were transformed into continuous time series. In Supplementary Figure 1, we illustrate the transformation where we convolve all spikes with two different kernels: a Gaussian kernel and an exponentially decaying kernel. The choice of kernel does not substantially alter the results presented here; therefore, we opted for the latter, as it prevents the introduction of current spiking information into the past. Additionally, the decay time was set to 5 ms to approximate the decay time of AMPA receptors, ensuring that the resulting output resembles the excitatory synaptic currents observed in downstream network layers.

#### Mutual Information

Mutual Information (MI) captures the dependence or shared information between two variables, providing insight into how much knowing one variable reduces uncertainty about the other. This metric is non-negative, with a value of zero indicating complete independence between the distributions of the two variables. The MI is calculated using the following expression:

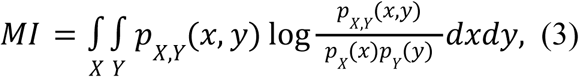

where X(t) and Y(t) are continuous random variables, with probability distributions p_X_ and p_Y_ and joint probability distribution p_X,Y_. Since the exact probability distributions are unknown and we only have a sample of them, estimating the distributions of Eq. 3 is not straightforward. To that matter, we utilized the “mutual_info_regression” function from scikit [44] which employs a nearest-neighbor approach for the estimation of the probability densities function. In cases where we calculate MI between the input and output of networks (e.g., Fig. 6c and 6d), the analysis was repeated while temporally shifting the ouput to account for delays in the integration. For example, the observed differences in the delay at which MI peaks between the encoding of the θ input (Fig. 6c) and the parallel pathway (Fig. 6d) primarily reflect their distinct, explicitly set, transmission delays: 20 ms for θ and 0 ms for the parallel pathway.

## Results

### Cross-frequency coupling and directionality depend on the connectivity

As an illustrative example and foundation for the present modeling work, we present electrophysiological recordings from the hippocampus of rats freely exploring known (familiar) and novel environments. These data revealed strong CFC between the phase of theta oscillations and the amplitude of gamma oscillations in pathway-specific local field potentials (LFPs) [12]. Figure 1a depicts the pathway-specific LFP corresponding to the CA1 *lacunosum moleculare* response to the input originating from the entorhinal cortex layer III (ECIII). The same study reported that both positive and negative CFD values coexist (Fig. 1a-iii); however, the predominant directionality, CFD_avg_, calculated as the average CFD across frequencies with significant CFC, was negative. This suggests that, on average, the amplitude of gamma oscillations preceeds the phase of theta. Following previous studies, all CFD values reported here were renormalized to emphasize regions of high CFC, with statistical significance (p < 0.01) determined through surrogate analysis of clusters with the highest absolute CFD values (for more detail see the analysis section and refs. [29,12].

To investigate theta-gamma interactions, we adapted the well-established circuit motifs Interneuron Network Gamma (ING) and Pyramidal-Interneuron Network Gamma (PING) [33, 35–37], with the addition of an external theta input (Fig. 1b-i and 1c-i). We began by analyzing separately the two circuit motifs, named as θ-ING and θ-PING respectively. Both models included a population of pyramidal cells (PCs) excited by a θ-modulated external input, with gamma rhythms generated locally by a population of fast-spiking self-inhibitory interneurons, the basket cells (BCs), which also projected to the soma of the PCs. All connections and parameters for the two motifs were identical, except for the nature of the BC-PC inhibition: feedforward in θ-ING and feedback in θ-PING. For more details on the model see methods section.

The distinct dynamics of these two circuit motifs are shown in Fig. 1b-ii and Fig. 1c-ii, where we depict the raster plots for the PCs and BCs (blue and orange dots, respectively). In the same panels we superimpose the mean somatic membrane potential (V_PC_) and the mean transmembrane somatic currents i_transm_, across all PCs. Both motifs exhibit strong theta-gamma CFC (see Fig. 1b-ii,vi and 2b-ii,vi) but, importantly, opposite directionalities were measured by CFD.

In the θ-ING model (see Fig. 1b-v,vi), gamma activity preceded theta oscillations, resulting in a negative CFD in areas of high CFC, similar to what was observed experimentally (see Fig. 1a). To investigate this further, we first ensured that the observed negative CFD was not an artifact of higher theta harmonics or non-sinusoidal waveforms, which could spuriously generate CFC and CFD values [45]. We systematically varied the transmission delay τ_θ,PC_ = 10/20/30 ms between the theta input and the PCs while keeping the transmission delay between the theta input and the BCs constant at τ_θ,BC_ = 20 ms. Thus we effectively varied the relative transmission delay Δτ = τ_θ,BC_ - τ_θ,PC_ that controls the temporal relationship between gamma and theta (Fig. 2). For Δτ = −10 ms, where the theta input reaches BCs earlier than the PC dendrites, gamma bursts occur at the rise of the theta oscillation, resulting in a more negative CFD (Fig. 2b, top panel). Conversely, for Δτ = 10 ms (Fig. 2b, bottom panel), gamma is relayed to PCs later, causing CFD to switch to positive. For comparison and completeness we also considered Δτ = 0 ms, in which theta input reaches both populations simultaneously, yet still results in a negative CFD (Fig. 2b, middle panel; same as in Fig. 1b). Overall, these results rule out a spurious contribution of theta harmonics or wave shape to the sign of CFD in the θ-ING motif.

To explain why gamma-to-theta interaction is detected when no relative transmission delay between them exists, we examined the relationship between the externally imposed theta input and the neuronal activity of both PC and BC populations (Fig. 2c and 2d red curve). To make this relation clearer, we represented PCs and BCs spikes firing relative to the phase of the theta input measured in the distal dendritic compartment of the PC (see Supplementary Fig. 1 for the same analysis using the external θ input as reference). When Δτ = 0 ms, BC activity increases faster than the external input to the PCs (Fig. 2c). Consequently, feedforward gamma inhibition reaches the PC soma before the excitatory theta activity propagates from the distal dendritic compartment. For completeness, we also include in Fig. 2c and 2d the cases for positive (in green) and negative (in orange) delays.

In contrast, in the θ-PING model, each cycle began with PC depolarization driven by the theta input, which, through the feedback connection, recruited BCs and induced gamma rhythmicity. Not surprisingly, in this configuration theta activity precedes gamma oscillations.

### Synaptic weights influence gamma-to-theta interactions in the θ-ING motif

We further explored the θ-ING results, since they better matched the experimental findings [12, 31]. To elucidate the synaptic mechanisms underlying gamma-to-theta interactions, we systematically increased the synaptic weights within the θ-ING model and evaluated their impact on CFC and neuronal theta firing patterns (Fig. 3). Notably, our findings indicate that CFD in the θ-ING model remains negative across a broad range of synaptic weights (see Table S3) and PC activity (see firing rates in Table S4), with positive or negligible CFD values observed only at extreme parameter values.

We first examined the influence of synapses projecting to PCs. When inhibition was very high relative to excitation (Fig. 3a-iv and 3b-i) PCs spiked only when the BCs’ activity was at its lowest value. Thus pyramidal spiking was scarce and out of phase with the external input, and the CFD was positive. Conversely, when excitatory input was excessively high relative to inhibition, PCs dominated the network dynamics, overriding BC-driven gamma oscillations (60–80 Hz), thereby disrupting CFC within this frequency band (Fig. 3a-i and 3b-iv). This effect was particularly pronounced when synaptic excitation surpassed the dendritic spiking threshold (Fig. 3b-iv). Under these conditions, the external theta generator retained the control over the network dynamics, with CFC shifting toward lower frequencies. In contrast, when excitation and inhibition were balanced, PC spiking occured slightly after the peak of the synaptic input (Fig. 3b-v). Then, the phase delay between input arrival and spiking was modulated by the level of the excitation/inhibition balance. Moreover, an increased inhibition broadened the phase distribution of PCs spiking and weakened the transmission of afferent theta signals to subsequent layers.

We next investigated the role of excitatory input to BCs (Fig. 3c). This synaptic connection not only regulated BC activity and, consequently, the total inhibitory drive to PCs, but also modulated the gamma frequency of the network. An increased excitatory drive to BCs accelerated the network dynamics, a result consistent with previous computational studies of ING models [46]. Overall, our results demonstrate that θ-ING networks exhibit flexible dynamics, capable of modulating both single-cell and population-level activity by changing synaptic weights (synaptic plasticity), while remaining in a mode where gamma leads theta locally.

### Circuits of combined θ-ING and θ-PING motifs

Given that brain circuits are generally endowed with both feedforward and feedback inhibition simultaneously, we next investigated the predominant theta-gamma directionality in the combined model, by adding the feedback PC→BC connection to the θ-ING model or, equivalently, the feedforward θ→BC connection to the θ-PING model. Under these conditions, we found that theta-gamma directionality was primarily determined by feedforward inhibition, as indicated by changes in CFD when the strength of the θ→BC connection is varied (Fig. 4a). In contrast, changing the strength of the feedback PC→BC connection resulted in significantly less pronounced changes (Fig. 4a).

**Fig. 4.**
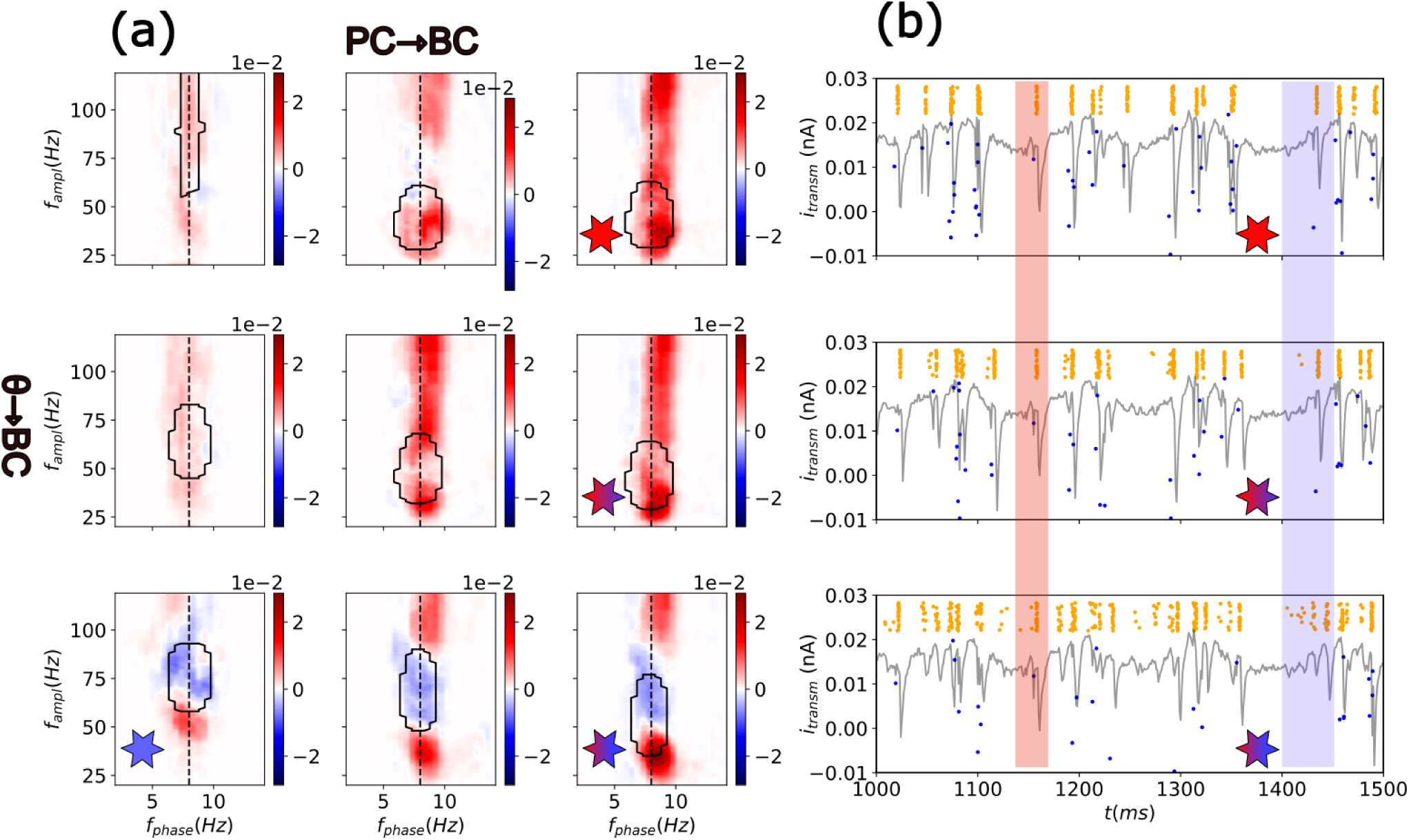
Transitions between positive and negative CFD in a combined θ-ING and θ-PING model. (a) CFD in the 2D parameter space of the relevant synaptic weights PC→BC and θ→BC (see Table S5). The θ→BC input strength increases in rows from top to bottom. The PC→BC input strength increases in columns from left to right. The red and blue stars denote the parameters of the pure θ-PING and θ-ING motifs, respectively (same motifs shown in Fig. 1). Stars of mixed colors depict the transition from a pure θ-PING to mixed models by increasing the feedforward connection strength θ→BC. (b) Transmembrane currents and raster plots of the θ→BC transition (same stars code as in panel a). Coloured rectangles highlight the initial part of a θ oscillation as well as the first few gamma cycles. The rectangle in blue depicts an example of directionality transition from top (CFD>0) to bottom (CFD<0). Note how increasing feedforward inhibition advances the firing of BCs (orange dots in the raster plot), and consequently the gamma oscillations over the theta cycle and the firing of PCs (blue dots). The rectangle in red depicts one example of theta cycle in which transition was not fully realized, highlighting the dynamic character of the CFD.

**Fig. 5.**
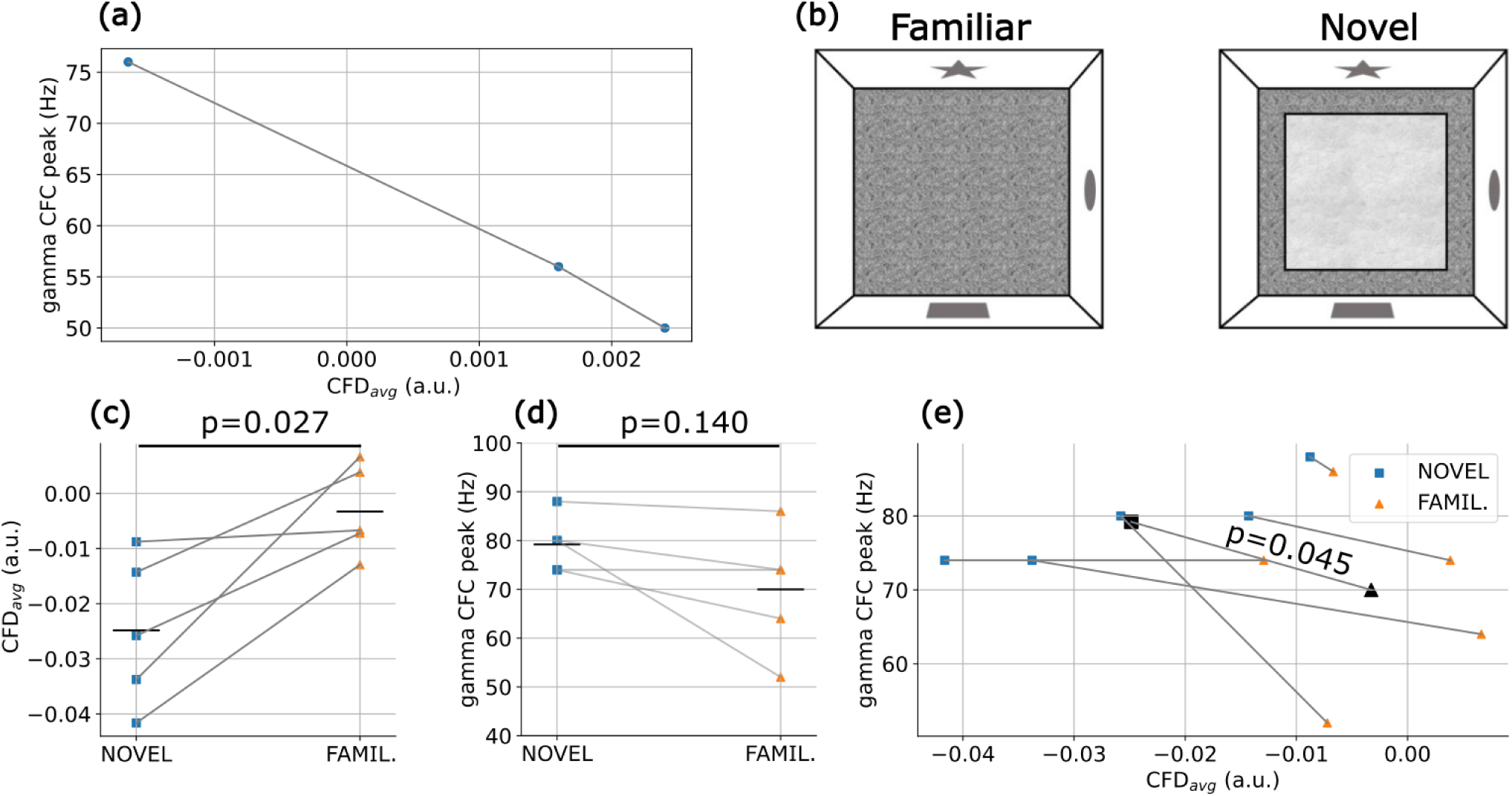
Experimental test of model predictions. (a) Mixed θ-ING/θ-PING motifs show that more negative CFD values are associated with higher gamma peak frequencies (same motifs as in the middle panel of Fig. 4a). (b) Schematic of the experimental setup: after one week of habituation with daily sessions in an open field, animals (n = 5) were introduced to a novel floor surface (mismatched novelty environment). Hippocampal recordings were performed during free exploration of both environments [12]. (c) CFD values were significantly lower during exploration of the novel environment compared to the familiar one (paired t-test t(4) = -3.40, p = 0.027). (d) During exposure to novelty, CFC tended to peak at higher gamma frequencies compared to the familiar condition, although this effect did not reach statistical significance (paired t-test t(4) = 1.84, p = 0.140). (e) Combined data from (c) and (d) represented in two dimensions. The slopes of all pairs of observations per animal (connected colored symbols) are significantly negative (one-tail t-test t(4) = -2.22, p = 0.045), demonstrating the association between CFD values and peak gamma frequencies. Black symbols represent average values across all rats for both conditions.

**Fig. 6.**
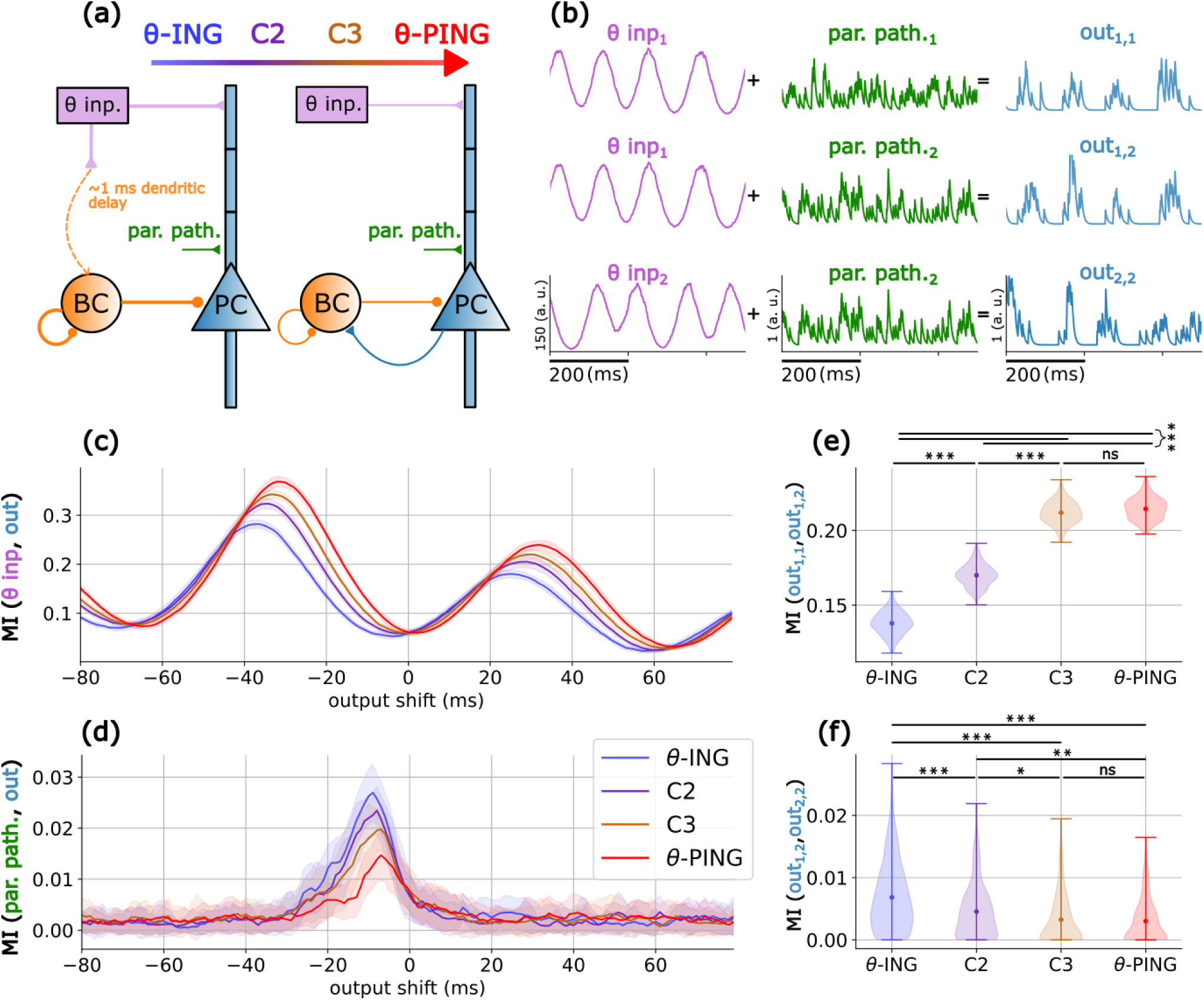
Functional analysis of θ-ING and θ-PING motifs. (a) θ-ING and θ-PING motifs with the parallel pathway introduced at the proximal dendritic segment of the PCs. Cases 2 and 3 (C2 and C3, respectively) are motifs in which the PC→BC and θ→BC synaptic weights fall between the values associated with the θ-ING and θ-PING motifs (see Table S6). (b) Schematic representation of the inputs in the θ (purple) and parallel (green) pathways and the outputs produced by the PCs (blue). All time series were generated by convolving the spike trains with a 5 ms exponential decay kernel. (c) Mutual Information (MI) between the θ input and the pyramidal output. Pairwise t-tests with Bonferroni correction for multiple comparisons between the maximum MI values showed that all cases are significantly different (p<0.001). (d) Same as in (c) but considering the parallel pathway as input. All cases are significantly different (p<0.001) except for C3 *vs.*θ-PING (p=0.0104), θ-ING *vs.* C2 (p=0.273) and C2 *vs.*C3 (p=0.103). (e) Consistency measured as MI between pairs of outputs in response to the same θ input while all other inputs changed. (f) Same as in (e) but considering the parallel pathway. Stars in panels (e) and (f) depict significant differences using pairwise t-tests with Bonferroni correction:***/**/* p<0.001,0.01,0.05 respectively.

To understand better the differential role of these connections, we focused on the dynamics of motifs transitioning from a pure θ-PING model to a mixed model with stronger feedforward connections (Fig. 4b). By increasing the external θ excitatory drive to BCs, this input overcomes the excitation received from PCs through the feedback PC→BC connection. Consequently, BCs fire progressively earlier relative to the theta phase, resulting in a predominant gamma-to-theta directionality (see blue rectangle in Fig. 4b). Simultaneously, the earlier inhibition of PCs further diminishes the influence of the feedback connection (PC→BC). Therefore, a unified model incorporating both feedforward and feedback connections suggests that CFD can be flexibly controlled via a local feedforward inhibitory connection.

### Evidence of θ-ING/θ-PING motifs in experimental data

The results obtained from the model provided at least two testable predictions. First, there is a relationship between CFD and the frequency of gamma oscillations coupled with theta, such that the more negative the CFD, the higher the gamma frequency (Fig. 5a). This relationship arises because in θ-PING, unlike in θ-ING, gamma generation depends on the spiking of PCs which are intrinsically slower than BCs. As the strength of the feedforward input to BCs increases, gamma generation becomes less dependent on PC firing, while its frequency progressively increases (Fig. 4a). Second, the quantitative CFD value depends on the balance between feedforward and feedback inhibitory pathways, making it potentially sensitive to different cognitive states.

Next, we sought to test these predictions using previously acquired electrophysiological data recorded from the hippocampus of 5 rats freely exploring familiar and novel spatial arenas (see Methods for details). We reasoned that exploring these distinct environments, which impose different cognitive demands—retrieving previously acquired spatial information versus encoding new information in a novel environment—would provide an effective testbed for identifying dynamic regulations of CFD. Indeed, this dataset revealed statistically significant differences in theta-gamma CFC [12].

We tested our predictions in the experimental dataset and found that, indeed, CFD was significantly modulated by the behavioural task, showing more negative CFD values during the exploration of the novel environment compared to the familiar one (Fig. 5b and 5c). In parallel, the gamma frequency exhibited a tendency to increase across all animals during the novelty condition (Fig. 5d), although this effect did not reach statistical significance (p = 0.13). However, we found a significant association between CFD and gamma frequency across all animals, consistent with model predictions, showing that more negative CFD values were linked to higher gamma frequencies in the hippocampus (Fig. 5e). Overall, these experimental findings support both predictions of the θ-ING/θ-PING framework.

### Functional differences of θ-ING and θ-PING motifs

Given the experimental evidence of the dynamic behaviour of cross frequency interactions, we explored whether different directionality modes, ranging from pure θ-ING to pure θ-PING, with two intermediate mixed motifs (Cases 2 and 3, Fig. 6a), exhibit distinct computational properties. Recognizing that in natural circuits, more than one pathway may converge into a neural population, we extended our analysis to include two external input pathways: (1) a θ-driven input targeting the distal dendrites of the PCs, as in previous analyses, and (2) a Poisson-distributed pulse input, termed the parallel pathway, targeting the proximal dendritic segment (see methods for more details).

To evaluate the computational properties of each motif, we employed two protocols. The first one assessed the encoding capacity by calculating the mutual information (MI) between an external input and its corresponding PC firing output. The second protocol evaluated the robustness of an input to variability. This is measured as the MI between two outputs generated in response to the same input, either the θ or the parallel pathway, while all other inputs varied (see Fig. 6b and the methods section for details on the protocols).

While all four motifs effectively encoded information from both the θ and parallel pathway inputs, they did so in a complementary manner. PING-shifted configurations were more effective at encoding the θ input, the primary driver of the circuit (Fig. 6c), whereas ING-shifted configurations more effectively encoded information from the parallel pathway (Fig. 6d). This disparity between motifs arises from a fundamental difference, evident in the membrane potential traces shown in Fig. 1b and 1c. In the pure θ-PING case, PCs and BCs are strongly coupled via feedback connections, resulting in larger gamma oscillations than in the θ-ING motif. This creates a robust theta-gamma scaffold where opportunity windows for encoding are restricted to the peaks of the gamma cycles nested within each theta cycle. Inputs that are not synchronized with this rhythmic theta-gamma structure, such as the parallel pathway, more often fail to evoke PCs responses. In contrast, in the θ-ING motif, gamma oscillations are primarily driven by BCs in a feedforward manner and, while still nested within the theta cycle, this gamma activity is more loosely anchored to PC firing, broadening the temporal windows for encoding. Consequently, strong perturbations falling within or outside the theta-gamma coding framework are able to overcome depolarization thresholds. To support this interpretation, we conducted simulations where the parallel pathway was replaced with single pulse perturbations. This allowed us to assign a gamma phase to each perturbation, revealing that the θ-PING motif was responsive during the peak of the gamma oscillation, whereas the θ-ING motif exhibited firing throughout the cycle (see Supplementary Fig. 3), confirming our hypothesis.

Finally, we assessed the second protocol, namely the robustness of an input to variability. Information encoding of the θ-input in PING-shifted motifs (Fig. 6c) was accompanied by greater consistency (Fig. 6e). Similarly, improved encoding of the parallel pathway in ING-shifted motifs (Fig. 6d) was also more robust to variability (Fig. 6f). Overall, the continuous transition observed between the two models, modulated by the balance between feedforward and feedback inhibition, suggests a mechanism for weighting or selecting communication channels with different temporal structures, which could be reflected in the CFD index.

## Discussion

In this study, we have shown that the directionality of theta-gamma interactions depends on, and can be regulated by, the feedforward and feedback inhibition balance. Specifically, we found that motifs with enhanced theta-modulated feedforward inhibition (θ-ING) exhibit dominant gamma-to-theta interactions, while those with enhanced theta-modulated feedback inhibition (θ-PING) display dominant theta-to-gamma interactions. In a combined circuit containing both feedforward and feedback connections, we found that the feedforward connection determines theta-gamma directionality and governs the transition between motifs or operational modes.

We further found that these operational modes significantly influence the firing phase of PCs within the theta cycle. As evident from Fig. 2d, the fine-tuning of gamma-theta interactions defines the temporal windows within the theta cycle in which cell assemblies can be formed. In this manner, the feedback-feedforward inhibitory balance may implement a push-pull mechanism that governs the firing phase of PCs: increased feedforward inhibition leads to phase precession and increased feedback inhibition induces phase recession. For the same reason, different operational modes favor the response of PCs to inputs with different temporal structures. By fine-tuning the timing between slow and fast oscillations, this mechanism selectively prioritises information transmitted through different afferent pathways, enhancing computational flexibility.

An additional notable observation was that the timing of theta-nested gamma inhibition modulated the phase difference between the afferent theta rhythm and the local theta spiking profile (Fig. 2d). This mechanism may facilitate the coordination of theta rhythms across pathway-specific field potentials, as observed experimentally [12]. Importantly, our analysis of hippocampal electrophysiological data supports this dynamical framework, demonstrating a behavioral state-dependent regulation of cross-frequency interaction directionality. In the following sections, we explore the practical applications and limitations of our model and discuss its relevance to experimentally observed brain dynamics.

### Is θ-ING a realistic model for negative CFD?

The emergence of negative cross-frequency directionality in the θ-ING motif is critically dependent on the rapid response of BCs, enabling gamma oscillations to precede local theta rhythms. This raises an important question regarding the physiological validity of the conditions assumed in our model. We argue that our implementation of BC activation times is, in fact, conservative.

First, dendritic transmission in BCs is notably rapid, with evoked postsynaptic currents at distal dendritic sites depolarising the soma within less than 1 ms dendritic delay [41]. Second, stimulation of the Schaffer collateral induces monosynaptic excitation followed by disynaptic inhibition after only 1.9 ± 0.2 ms delay [47]. This is faster, on average, than the delays assumed in our model, which consists of the 1ms dendritic delay plus 1.5 ± 0.2 ms of synaptic delay from BC to PCs and the spike generation time. Third, both *in vivo* and *in vitro* patch-clamp recordings from the CA1 region have demonstrated faster action potential initiation in BCs compared to PCs following stimulation of pathways that simultaneously excite BCs and PCs, namely the perforant path and the Schaffer collateral, respectively [48, 49]. Finally, regarding population dynamics, *in vivo* recordings from mice running in a maze have shown that interneurons activity in the pyramidal layer of CA1 peaks approximately 60 degrees (or equivalently 20 ms) ahead of the theta phase of PCs (see Fig. 5 in ref. [50]). This finding is consistent with our results (Fig. 2c and 2d), which show that BCs lead PCs significantly in theta phase. Taken together, these findings support the plausibility of the inhibitory time delays implemented in our model.

### Experimental evidence and interpretation of CFD measurements

Our results indicate that the balance between feedforward and feedback inhibition in local circuits determines the directionality of cross-frequency interactions. We further showed that a feedback-shifted balance favours transmission in an afferent pathway by promoting the specific cross-frequency rhythmicity driven by the afferent input, while a feedforward-shifted motif broadens the opportunity window for encoding, facilitating parallel pathways to transmit. Accordingly, dynamic CFD measures could be interpreted in terms of predominant inhibitory circuit motifs and flexible prioritization of functional connectivity pathways.

In the rat hippocampus, we have found that during novelty exploration, activity in the EC→CA1 connection (temporoammonic pathway) exhibits more negative CFD values than during exploration of familiar environments. According to our model, a less negative CFD corresponds to a PING-shifted state that favors transmission along the main afferent pathway (here the EC→CA1 connection) while decreasing competition from parallel pathways, such as the Schaffer collateral (driven by the trisynaptic DG→CA3→CA1 connections). In contrast, a more negative CFD suggests an ING-shifted state, where parallel inputs gain greater relevance. This interpretation aligns with previous findings from c-Fos interaction networks, which reported an activity shift from a dominant temporoammonic monosynaptic pathway during familiar conditions to a dominant trisynaptic pathway during novelty exploration [51]. Animals in the mismatch novelty condition compare the contextual representation retrieved from memory with incoming sensory information. Consistent with established perspectives on information processing within the hippocampal circuit [52, 53, 22, 12], our model results indicate that novelty conditions favor an ING-shifted operational mode. This mode facilitates the integration of parallel pathways, enabling the retrieval of memories (CA3→CA1) while simultaneously processing environmental cues and encoding new information (EC3→CA1).

Furthermore, a recent study using electrocorticography in human epilepsy patients performing a spatial attention task reported a relationship between alpha-gamma CFD values and attentional states [30]. Specifically, more negative CFD values were associated with non-attended stimulus, while lower absolute (but still negative) CFD values were linked to attended stimulus. According to our model, less negative CFD would be reflecting a PING-shifted circuit state that favors transmission of the message in the afferent (attended) pathway, in this case, the visual dorsal stream, reducing the impact of parallel (distracting) inputs. Similarly, a more negative CFD, as found for non-attended stimulus, is expected from an ING-shifted circuit in which parallel inputs gain relevance over the afferent pathway. Importantly, the authors reported enhanced functional connectivity along the visual dorsal stream in the first case, and suppressed connectivity in the second, supporting our interpretation. The model offers a mechanism to explain attention deployment dynamically and flexibly, based on feedforward-feedback inhibitory balance and reflected in the CFD metric.

Interestingly, these studies also confirmed key mechanistic predictions of our theoretical framework. First, in the electrocorticography study alpha-phase activity in upstream regions preceded downstream high gamma activity, while locally, gamma preceded alpha, which aligns well with our model predictions (compare Fig. S2 with Fig. 2 in our manuscript and Fig. 2 of [30]). Second, both studies provide preliminary evidence supporting the mechanism of directionality control based on the feedforward and feedback inhibitory balance, showing that more negative CFD values are associated with higher peak gamma frequencies, as expected from the feedforward recruitment of BCs (Fig. 4 in our manuscript and Fig. 2 in [30]).

### Limitations

Our model is intentionally minimal, designed to ensure the reported directionalities are broadly applicable across different brain regions, as supported by existing literature [29, 31, 54, 55, 12, 56, 30]. This generalizability, however, comes at the expense of region-specific precision. While addressing such specificity would require detailed multicompartmental models tailored to particular brain areas [57, 58, 59], the simplicity of our model allows it to capture fundamental dynamics effectively. Future work incorporating diverse interneuron types and their unique connectivity patterns, particularly in regions like the hippocampus, would further refine our understanding [60].

While the motifs reproduce gamma-to-theta and theta-to-gamma directionalities, they do not account for large cross-frequency lags, such as the -50 ms lag between alpha and beta rhythms observed in human auditory cortex electrocorticogram data [31]. However, by accounting for plausible relative transmission delays, as illustrated in Fig. 2, the motifs can accommodate a wide range of experimentally observed cross-frequency lags. These lags may reflect multi-synaptic pathways. For instance, in the CA3 region, long-lag negative CFD could arise if BCs are monosynaptically recruited by entorhinal cortex inputs, while PCs are driven through the disynaptic circuit (EC→DG→CA3). Investigating these delays in specific anatomical contexts offers promising opportunities for future research.

We did not address different potential mechanisms of CFC generation. In our model, external theta rhythms drive local interneurons and PCs to generate local gamma and theta oscillations. This design aligns with patch-clamp recordings highlighting the role of local inhibition in gamma generation, as well as with LFP recordings between distant sites in the hippocampus that exhibit high coherence in the theta but not in the gamma band [26, 24, 61, 25]. However, in experiments recording theta-gamma CFC in connected layers of the hippocampal formation, it has been shown that the upstream theta-gamma activity may influence downstream CFC dynamics [21–23, 62]. These findings suggest that the generation of CFC may be contributed by both local and network mechanisms. Interestingly, in our model, the firing of the PCs, both in the θ-ING and θ-PING motifs, relays a theta-gamma coupled signal to a potential downstream target (see the CFC in the PCs population firing in Supplementary Fig. 1). Therefore, although our primary focus was on the directionality of cross frequency interactions, the model captures key insights into the generation of CFC and provides a valuable framework for further exploration. How do local and network mechanisms of CFC generation interact and how the interaction conditions firing sequences and information transmission? Future work will address this question.

In summary, despite its simplifications, the model offers a robust framework for understanding CFC and CFD across regions while paving the way for more detailed, region-specific explorations in future studies.

### Concluding remarks

In conclusion, our combined modeling and experimental results propose a mechanism for the flexible gamma-to-theta interactions observed in electrophysiological recordings. They support the view that both θ-ING/θ-PING operational modes exist along a continuum rather than as mutually exclusive alternatives and suggest a functional role for feedforward/feedback inhibitory balance in prioritizing parallel information pathways converging onto the same dendritic tree. Thus, CFD and related measures may serve as valuable experimental entry points for further elucidating these processes.

## Author Contributions

- Conceptualization: Dimitrios Chalkiadakis, Santiago Canals, Claudio R. Mirasso
- Model development: Dimitrios Chalkiadakis, Jaime Sánchez-Claros
- Analysis: Dimitrios Chalkiadakis, Santiago Canals, Victor J. López-Madrona
- Preparation of the manuscript: Dimitrios Chalkiadakis, Santiago Canals, Claudio R. Mirasso, Jaime Sánchez-Claros, Victor J. López-Madrona

## Acknowledgements

D. Chalkiadakis, J. Sanchéz- Claros and C. R. Mirasso acknowledge support from the Spanish Ministerio de Ciencia, Innovación y Universidades through projects PID2021-128158NB-C22 and María de Maeztu CEX2021-001164-M funded by the MICIU/AEI/10.13039/501100011033. D. Chalkiadakis and S. Canals acknowledge support from the Spanish Ministerio de Ciencia, Innovación y Universidades through projects PID2021-128158NB-C21 and Severo Ochoa CEX2021-001165-S funded by the MICIU/AEI/10.13039/501100011033

## Data availability statements

The electrophysiological datasets used in this study are available at: https://doi.org/10.20350/digitalCSIC/12537. Codes can be assessed at https://github.com/gerompampastrumf/thetaING_PING

## Supporting Information

**Table S1.** Synaptic parameters of the model. NMDA has an additional scaling factor due to the magnesium block 1/(1 + 0. 28[*Mg*]*e*-^0.062*V*^), where [Mg] = 1mM is the concentration of magnesium, and V the membrane potential in mV.

| Receptor | $\tau_r(\text{ms})$ | $\tau_d(\text{ms})$ |
| --- | --- | --- |
| AMPA | 0.05 | 5.3 |
| NMDA | 15 | 150 |
| GABA | 0.07 | 9.10 |

**Table S2.**
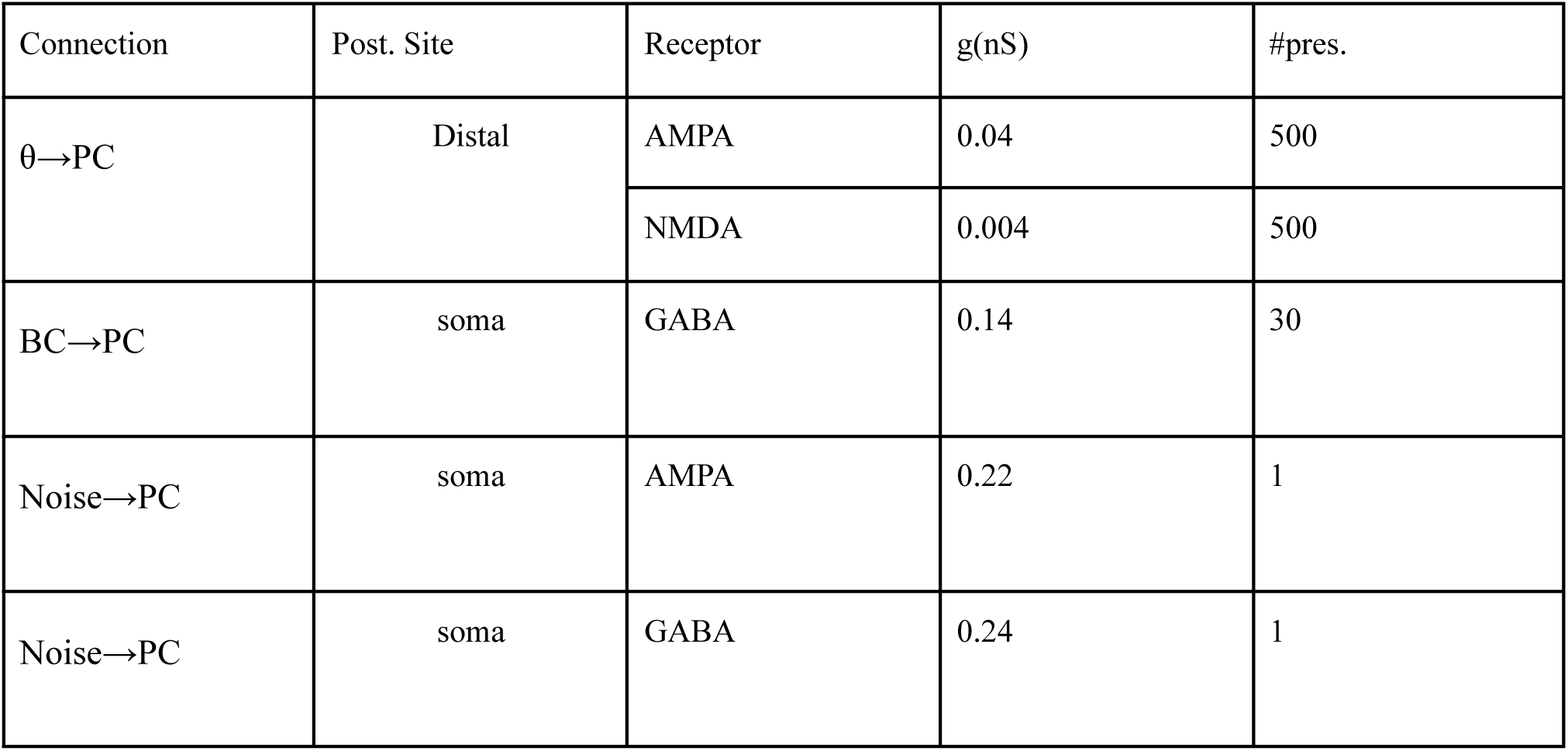
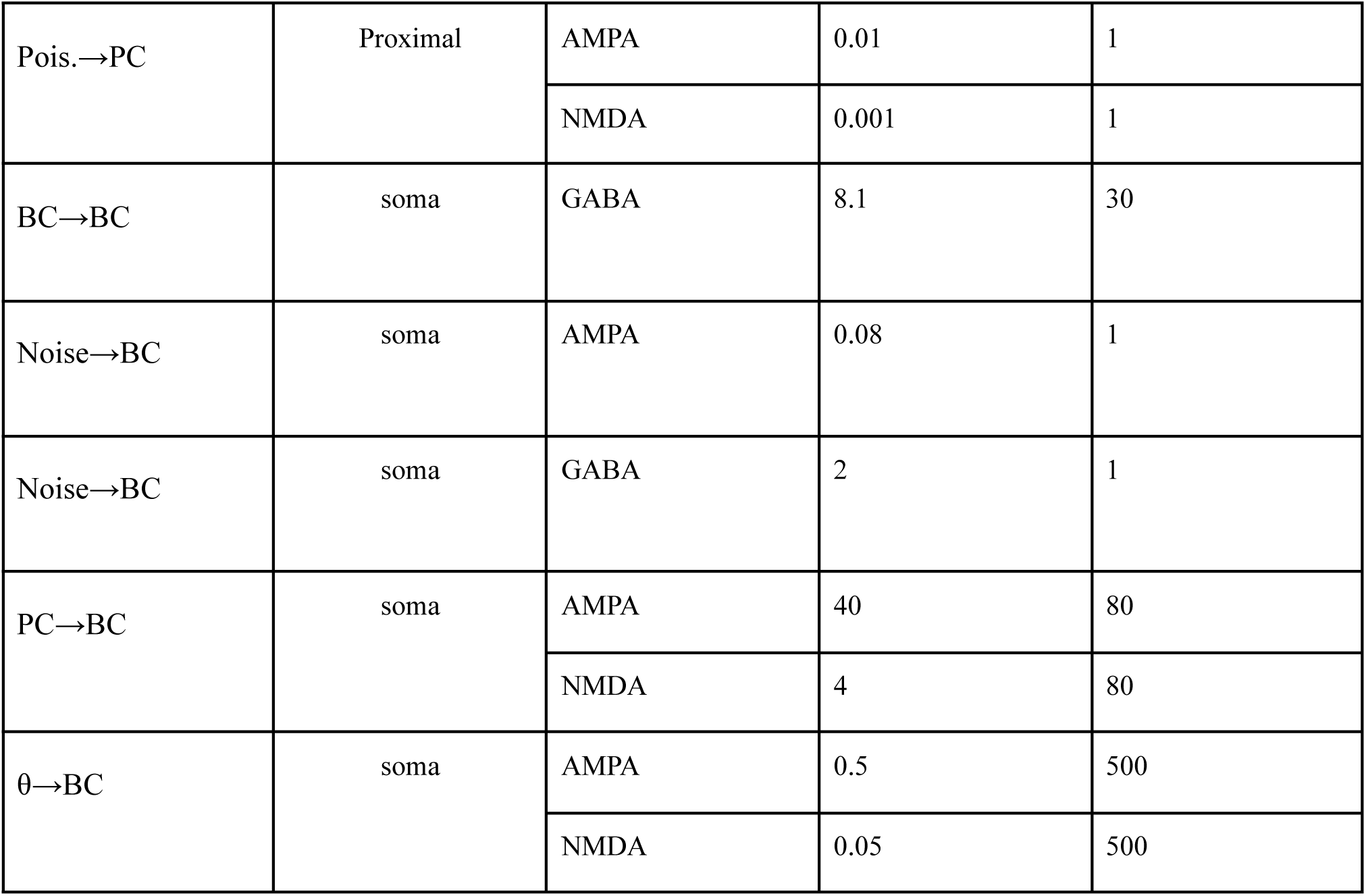
Synaptic weights for Fig. 1. The synaptic weight depicted for PC→BC is for the θ-PING, otherwise it is zero. The synaptic weight depicted for θ-ING is for the θ→BC, otherwise it is zero. The difference in the order of magnitude between PC→BC and θ→BC is due to differences in the number of presynaptic neurons (80 vs 500) and their activity (0.49Hz vs 8Hz).

| Connection | Post. Site | Receptor | $g(\text{nS})$ | #pres. |
| --- | --- | --- | --- | --- |
| $\theta$→PC | Distal | AMPA | 0.04 | 500 |
|  |  | NMDA | 0.004 | 500 |
| BC→PC | soma | GABA | 0.14 | 30 |
| Noise→PC | soma | AMPA | 0.22 | 1 |
| Noise→PC | soma | GABA | 0.24 | 1 |
| Pois.→PC | Proximal | AMPA | 0.01 | 1 |
|  |  | NMDA | 0.001 | 1 |
| BC→BC | soma | GABA | 8.1 | 30 |
| Noise→BC | soma | AMPA | 0.08 | 1 |
| Noise→BC | soma | GABA | 2 | 1 |
| PC→BC | soma | AMPA | 40 | 80 |
|  |  | NMDA | 4 | 80 |
| $\theta$→BC | soma | AMPA | 0.5 | 500 |
|  |  | NMDA | 0.05 | 500 |

**S1 Fig.**
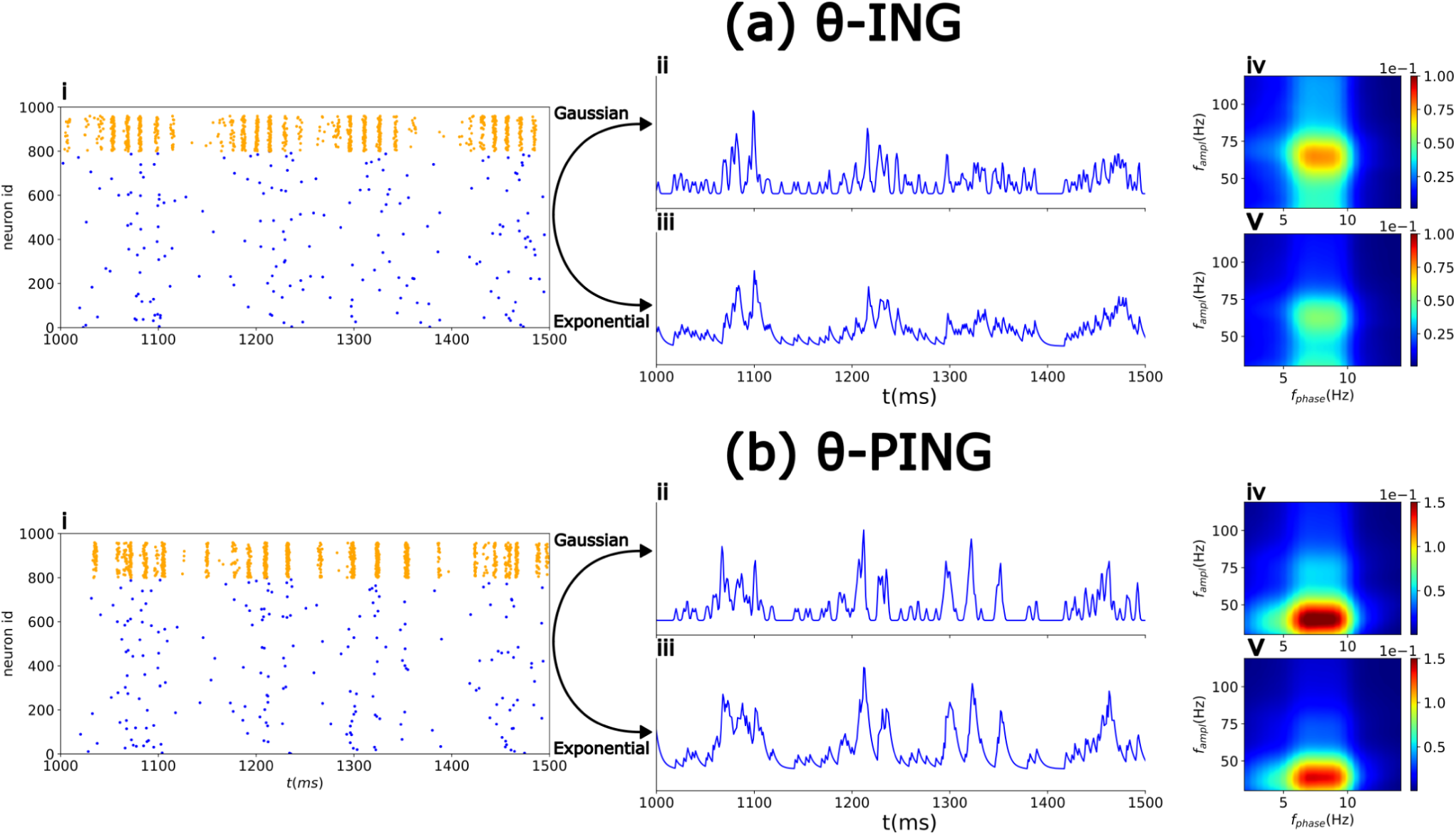
Generation of the PC output of the network. (i) Raster plots of PCs (and BCs for completeness). The convolution of each PC spike with either a 1 ms Gaussian kernel (ii) or a 5 ms decaying exponential kernel (iii) generates a proxy for the instantaneous output firing rate (in arbitrary units). A CFC analysis of the firing rate (iv, v) provides insight into what could be detected in the dendrites of a downstream layer which is not dependent on the kernel. This analysis was conducted for PC spikes in a θ-ING motif (a) and a θ-PING motif (b). To ensure sufficient spikes for the CFC analysis, both networks were scaled up by a factor of 4—achieved by increasing the number of neurons and projections while reducing the number of synaptic weights.

**S2 Fig.**
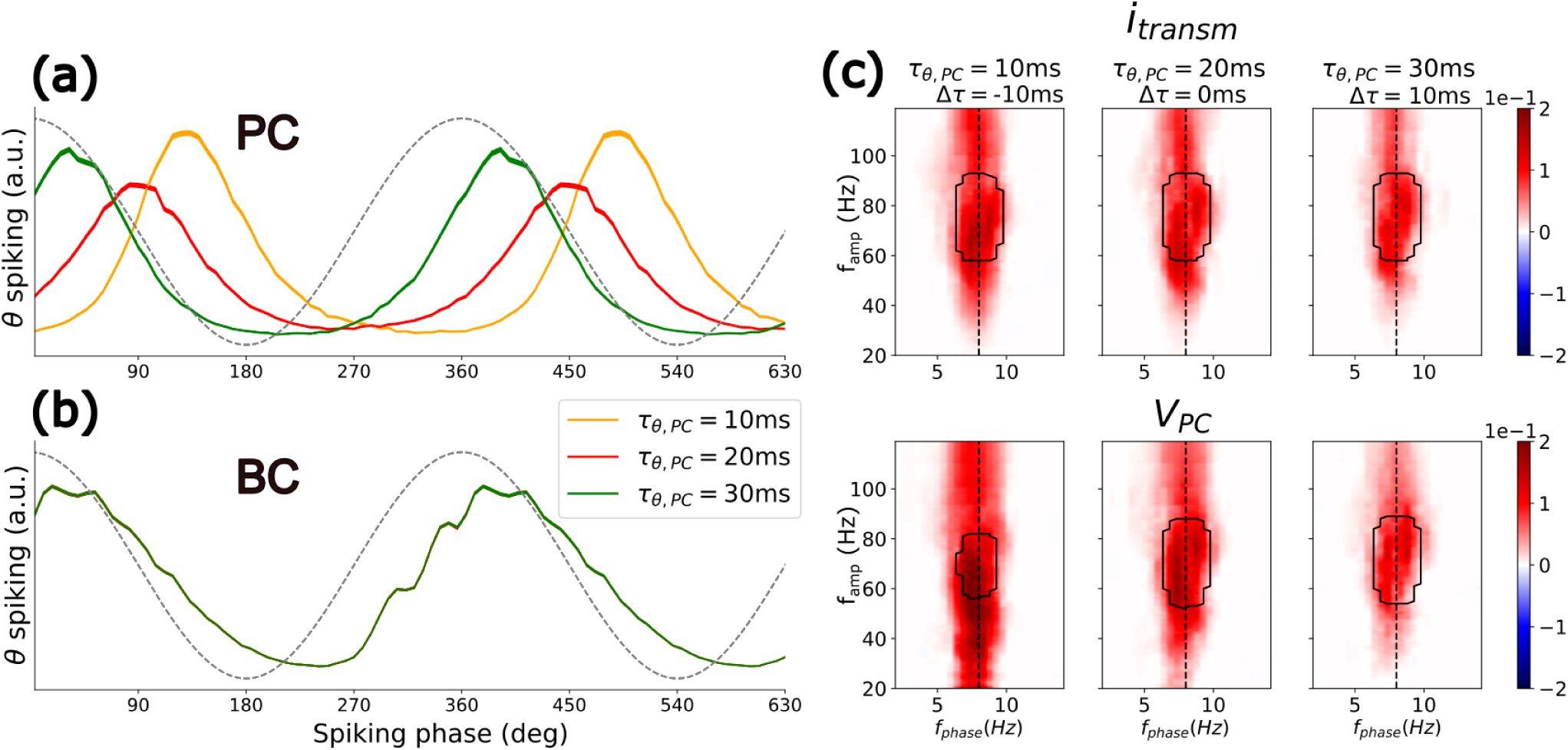
Similar to Fig. 2, but using the external θ drive phase as the θ reference for spiking and CFD calculations. Panels (a) and (b) show the θ phase of PC and BC spiking, respectively. Panel (c) illustrates the CFD for i_transm_ (top) and V_PC_ (bottom). To derive the θ phase of the external population, spikes are passed through a decaying exponential kernel with a 5 ms time constant. Note that when using the external population’s θ phase, the BC phase remains unchanged, while the PC phase shifts significantly due to different offsets. Finally, as the local γ is consistently generated by the external θ driver, the CFD remains positive under all conditions.

**Table S3.**
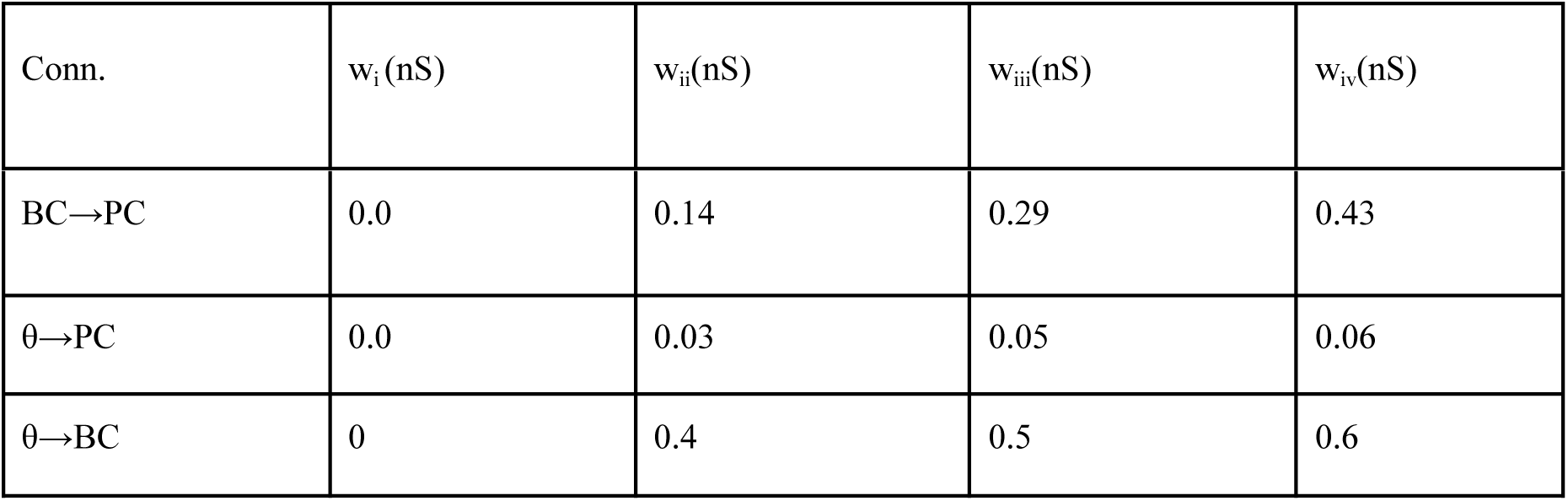
Synaptic weights for Fig. 3. All other synaptic parameters are as in θ-ING (see Table S2).

**Table S4.**
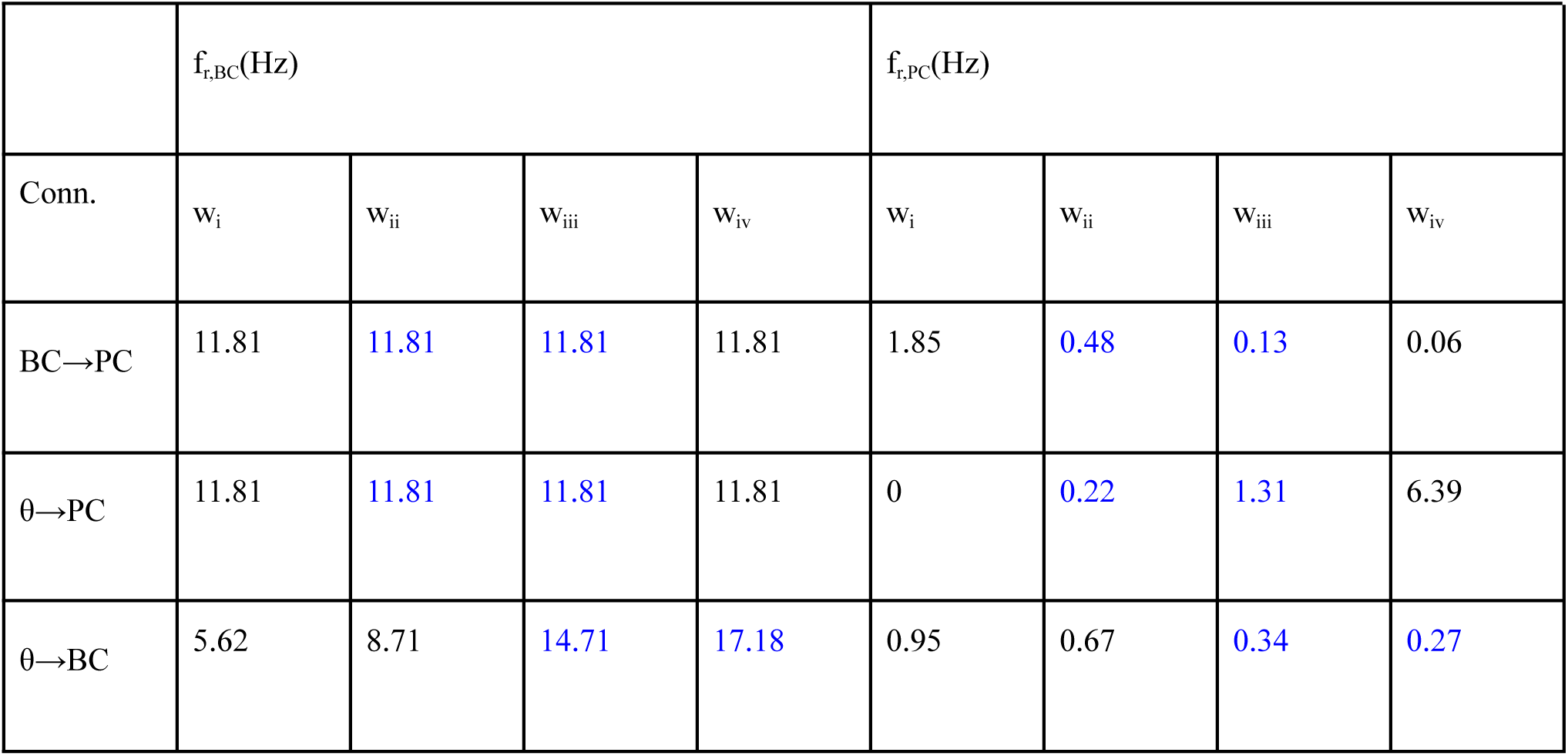
Mean firing rates for BCs (f_r,BC_(Hz)) and PCs (f_r,PC_(Hz)) in the θ-ING model for different synaptic weights **v =** (w_i_<w_ii_<w_iii_<w_iv_) in the circuit connection (Conn.). The simulations are the same as in Fig. 3. Blue-colored cells depict motifs that exhibit negative CFD.

**Table S5.** Synaptic weights for Fig. 4. As also stated in Table S2, the difference in the order of magnitude between PC→BC and θ→BC is due to differences in the number of presynaptic neurons (80 vs 500) and their activity (0.49Hz vs 8Hz).

| Conn. | $w_i$ (nS) | $w_{ii}$ (nS) | $w_{iii}$ (nS) |
| --- | --- | --- | --- |
| PC→BC | 0 | 20 | 40 |
| $\theta$→BC | 0.0 | 0.25 | 0.5 |

**Table S6.** Synaptic weights for Fig. 6.

| Conn. | θ-ING | C1 | C2 | θ-PING |
| --- | --- | --- | --- | --- |
| PC→BC | 0 | 10 | 20 | 60 |
| $\theta \rightarrow$BC | 0.5 | 0.42 | 0.33 | 0.0 |

**S3 Fig.**
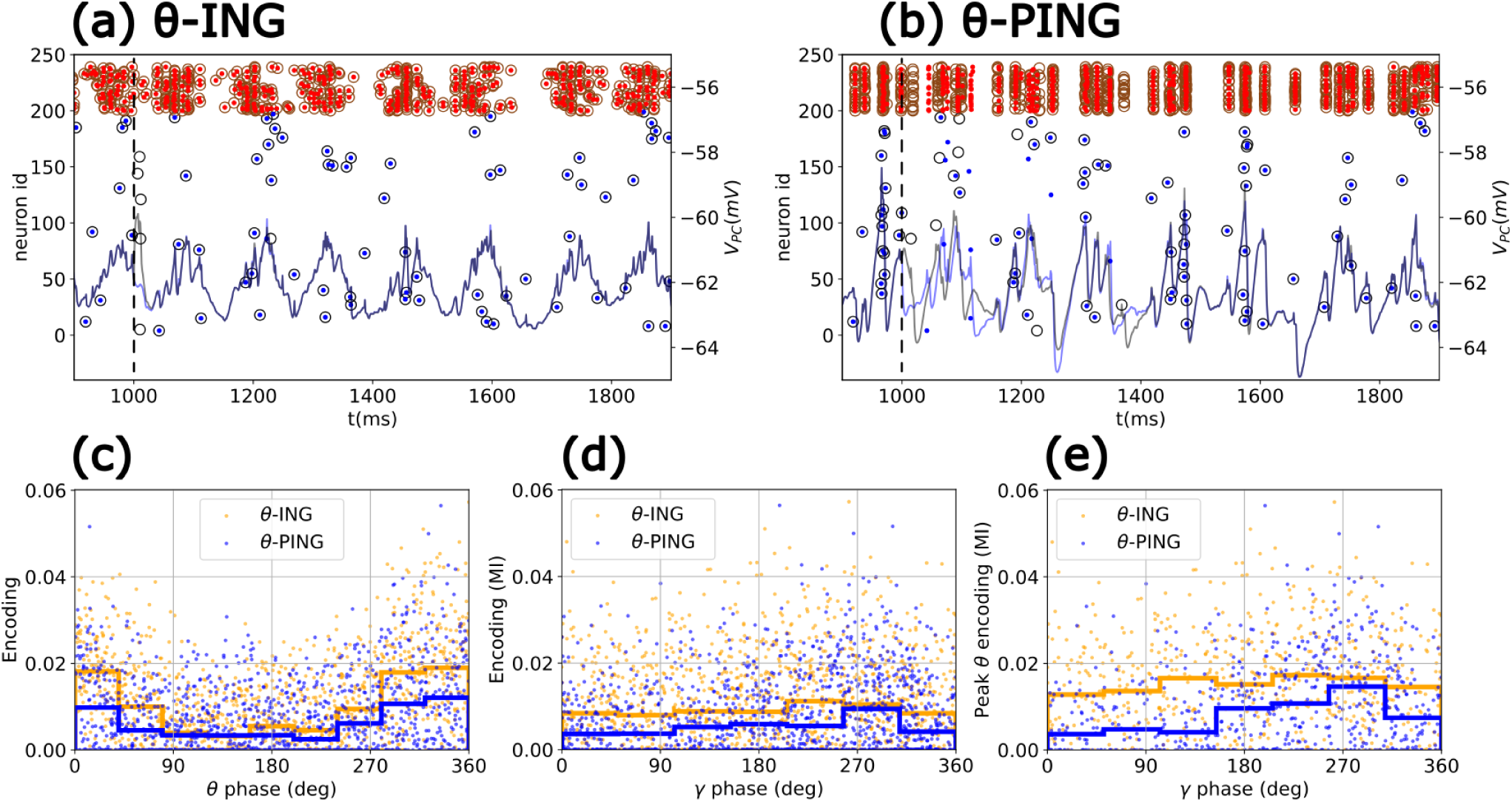
Single perturbation MI analysis. A single spike is introduced in the proximal dendrite at a predefined time within the interval (1-1.5) s (specifically at 1 s, indicated by the black dashed line). Panels (a) and (b) show the dynamics for a θ-ING and a θ-PING motif, respectively, both with similar firing rates. Blue lines represent the mean membrane potential at the PC soma (V_PC,g_) for the unperturbed case, while gray lines show the evolution after the perturbation (V_PC,p_). Open circles indicate spikes in the perturbed simulations (gray for PCs, brown for BCs), and solid circles represent spikes in the baseline condition (blue for PCs, red for BCs). Since only one perturbation is applied, the resulting encoding value between output and perturbation can be related to the network state at the time of perturbation. (c) Encoding values are plotted against the θ phase of V_PC,p_, where 180° represents the trough and 0°/360° the peak. (d) Same as (c), using a high-pass filter cutting of frequencies lower than 20 Hz to capture the gamma activity of both motifs. (e) Same as (d), but for encoding values only when the θ phase is between -90° and 90°, i.e., when the network is more depolarized.

